# Air-seq: Measuring air metagenomic diversity in an agricultural ecosystem

**DOI:** 10.1101/2022.12.13.520298

**Authors:** Michael Giolai, Walter Verweij, Neil Pearson, Paul Nicholson, Richard M. Leggett, Matthew D. Clark

## Abstract

**Background:** All species shed DNA into their environment during life or in death providing an opportunity to monitor biodiversity via its environmental DNA. Biodiversity monitoring using environmental DNA based technologies has become an important tool in understanding ecosystems. In recent years promising progress for non-invasive and, more importantly, non-destructive monitoring has been made by combining the retrieval of information transmitted by released environmental DNA with high-throughput sequencing technologies. Important ecosystems under continuous threat by disease but essential for food supplies are agricultural systems, often farmed as large monocultures and so highly vulnerable to disease outbreaks. Pest and pathogen monitoring in agricultural ecosystems is therefore key for efficient and early disease prevention and management. Air is rich in biodiversity, but has the lowest DNA concentration of all environmental media and yet it is required for windborne spread by many of the world’s most damaging crop pathogens. Our work and recent research suggests that ecosystems can be monitored efficiently using airborne nucleic acid information.

**Results:** Here we show that the airborne DNA of microbes can be recovered, sequenced and taxonomically classified, including down to the species level. Monitoring a field growing key crops we show that Air-seq can identify the presence of agriculturally significant pathogens and quantify their changing abundance over a period of 1.5 months often correlating with weather variables.

**Conclusion:** We add to the evidence that aerial environmental DNA can be used as a source for biomonitoring in agricultural and more general terrestrial ecosystems. The ability to detect fluxes and occurrence patterns of species and strains with high throughput sample processing and analysis technologies highlights the value of airborne environmental DNA in monitoring biodiversity changes and tracking of taxa of human interest or concern.

## Introduction

Air is rich in biodiversity containing particles of prokaryotic and eukaryotic organisms such as pollen, matter from animals and a variety of bacterial and fungal species [1]. Wind patterns are important in influencing climate [2], species dispersal, species biogeography [3–5] and plant as well as animal health [6, 7]. Especially when anthropogenic actions cause dramatic effects on the environment, measures to understand processes and species fluxes influencing ecosystems become of increasing importance [8]. One means to help characterise such changes is the utilisation of nucleic acids found as environmental DNA (eDNA) [9]. eDNA studies have become increasingly valuable in combination with high-throughput sequencing technologies [3]: for example searching for the term ‘environmental DNA° on PubMed highlights 72,257 published articles in the 16 years between 2005 (the year that pyrosequencing as an early high throughput sequencing technology debuted [10]) to 2021 in contrast to 18,782 results published in the 65 preceding years between 1959 (the start of PubMed records) to 2004. Hence, eDNA studies are especially helpful as they can be utilised to increase sample throughput at a reduced cost and facilitate profiling of environments in a non-destructive, non-invasive manner [9, 11].

Plant agricultural ecosystems are, as a source of food, essential for human needs [12]. Because modern farming is mostly practised in large monocultures [13, 14], disease - if not managed effectively - can have devastating effects on yield and food security [15–17]. Disease propagation and outbreaks are not solely based on close distance transmission of pathogens to neighbouring plants, but also from long distance aerial dispersal of pathogenic material such as fungal spores between fields [4]. Disease monitoring is therefore crucial to prevent and mediate pathogen damage [18]. In large patches of farmland this can be achieved with a multitude of technologies ranging from chemical, molecular to IoT (Internet of Things - i.e. systems of interconnected digital devices) technologies [19]. However, many current pathogen detection technologies rely on the detection and analysis of visual plant damage [19]. Highly specific molecular tools (e.g. PCR [20] or antibody-based assays [21]) can test for single pathogens but fail to distinguish between different strains and genotypes. Next generation sequencing based approaches are of great potential to overcome these limitations providing the means for pathogen detection before disease establishment at a resolution that can distinguish between pathogen strains.

One promising approach to monitor disease onset in agroecosystems is to monitor the species composition of air [6]. Air has already been a regular subject of study in the past to understand the species dynamics and composition of ecosystems (e.g. pollen dispersal) [1, 6, 7, 22–29]. eDNA sampled from air in combination with high-throughput metagenomic sequencing therefore can be a useful tool in profiling ecosystems. Many of the present studies however rely on selective and targeted metabarcoding strategies [1, 6, 7, 24, 25, 27, 29], while shotgun metagenomics is still challenging to achieve with low concentration DNA samples and, so far, very recently reported results are limited to urban areas [23, 26].

For example, a recent study conducted by Aguayo et al. [6] reported that the ash dieback causing fungus *Hymenoscyphus fraxineus* is present at higher frequencies in air samples of forests with confirmed ash dieback disease. Two other studies by Lynggaard et al. [29] and Clarke et al. [30] showed the high accuracy of air eDNA profiling in systems with described species. We hypothesised that monitoring air metagenomes in agricultural ecosystems will show a rich dynamic of pests and pathogens associated with the cultivated crops and will be able to identify newly appearing and emerging diseases over time. To test our hypothesis we established a custom workflow hereafter termed ‘Air-seq° utilising a field-work compatible, small, battery powered and portable air-sampling device (i.e. the Coriolis Micro Microbial Air Sampler) and a next-generation sequencing workflow preparing aerial eDNA shotgun metagenome sequencing libraries (**Figure 1A**). Using Air-seq we show the metagenome dynamics in the months of June and July in a field growing wheat barley and pea close to the Earlham Institute at the Norwich Research Park in Norwich, United Kingdom (UK) (**Additional file 1.docx**).

**Figure 1.**
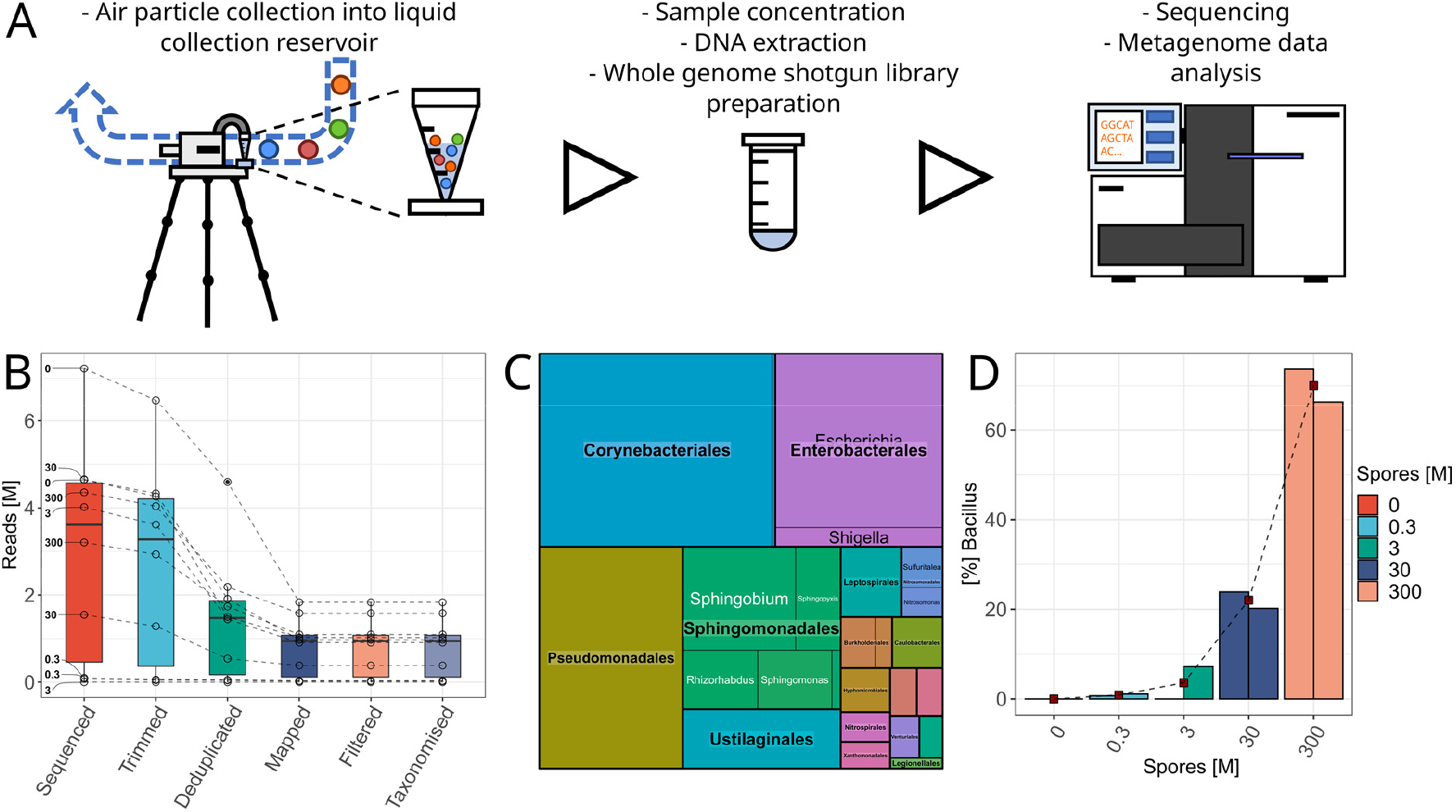
Air-seq performance in a wind-tunnel experiment while releasing *B. thuringiensis* spores. **(A)** Air-seq workflow: Air is filtered for a defined time-period through a liquid collection buffer reservoir. After sampling the liquid collection buffer is concentrated, the eDNA extracted from the sample and a whole genome shotgun library prepared using a low-input DNA compatible Illumina Nextera protocol. After library preparation the sample is ready for sequencing and data analysis. **(B)** Read numbers (million) over all analysis steps, from number of sequenced, trimmed, deduplicated, diamond aligned, quality filtered and taxonomically assigned reads. The spore concentration in million released spores is shown as the bold number to the left of the ‘Sequenced’ boxplot. **(C)** The background metagenome of the wind-tunnel; the larger the box, the more normalised reads of a specific order were detected; each order is grouped in boxes with genera, with the box size directly proportional to the abundance of the corresponding genus. **(D)** Detection curve of percent *Bacillus* in the classified reads of each spore concentration. For each spore concentration two replicates were sampled. When releasing only water (0 million spores) we could not detect any *Bacillus* calls. With an increase in the released spore number the abundance of *Bacillus* in the libraries rose rapidly.

## Results

### Detection of *Bacillus thuringiensis* spores in a controlled wind tunnel experiment

We first tested the accuracy and quantitative ability of our Air-seq method in a wind-tunnel experiment releasing a specific amount of *B. thuringiensis* spores: we continuously sprayed a total number of 0 (i.e. water as a background control), 0.3, 30 and 300 million spores over a time-period of 10 minutes at a distance of 5 m from the air sampler. For each spore concentration we collected two replicates with 10 minutes sampling time. For each replicate we prepared an Illumina Nextera sequencing library and sequenced the libraries to an average of 3.60 ± 1.65 million 150 base-pair single-end reads (**Figure 1B**). In the background control we observed *Corynebacteriales* (27.10 % of all counts), *Pseudomonadales* (19.10 % of all counts) and *Enterobacterales* (17.58 % of all counts) as highest abundant orders, with *Mycobacterium, Pseudomonas* and *Escherichia* as highest abundant genus in each order (**Figure 1C**). Releasing increasing amounts of spores, we observed a quantitative increase of the genus *Bacillus* in the samples (Pearson correlation coefficient: 0.97, R-squared: 0.95): 0 % of reads were assigned as *Bacillus* with releasing 0 million spores, 0.88 % of reads with 0.3 million spores, 3.59 % of reads with 3 million spores, 22.02 % of reads with 30 million spores and 70.04 % of reads with 300 million spores (**Figure 1D**).

### Determination of a suitable air sampling time-window for in-field metagenome measurements

After wind-tunnel experimentation we tested air collection in the field. We prepared a collection time-series sampling air for 5, 10, 30, 60 and 120 minutes consecutively on the same day at a Norwich Research Park field site close to the Earlham Institute (52°37’21.1”N, 1°13’02.6”E). For each time-point we collected two samples. We prepared Illumina Nextera sequencing libraries from the samples and sequenced the libraries to 5.14 ± 1.94 million 150 base-pair single-end reads (**Figure 2A**).

**Figure 2.**
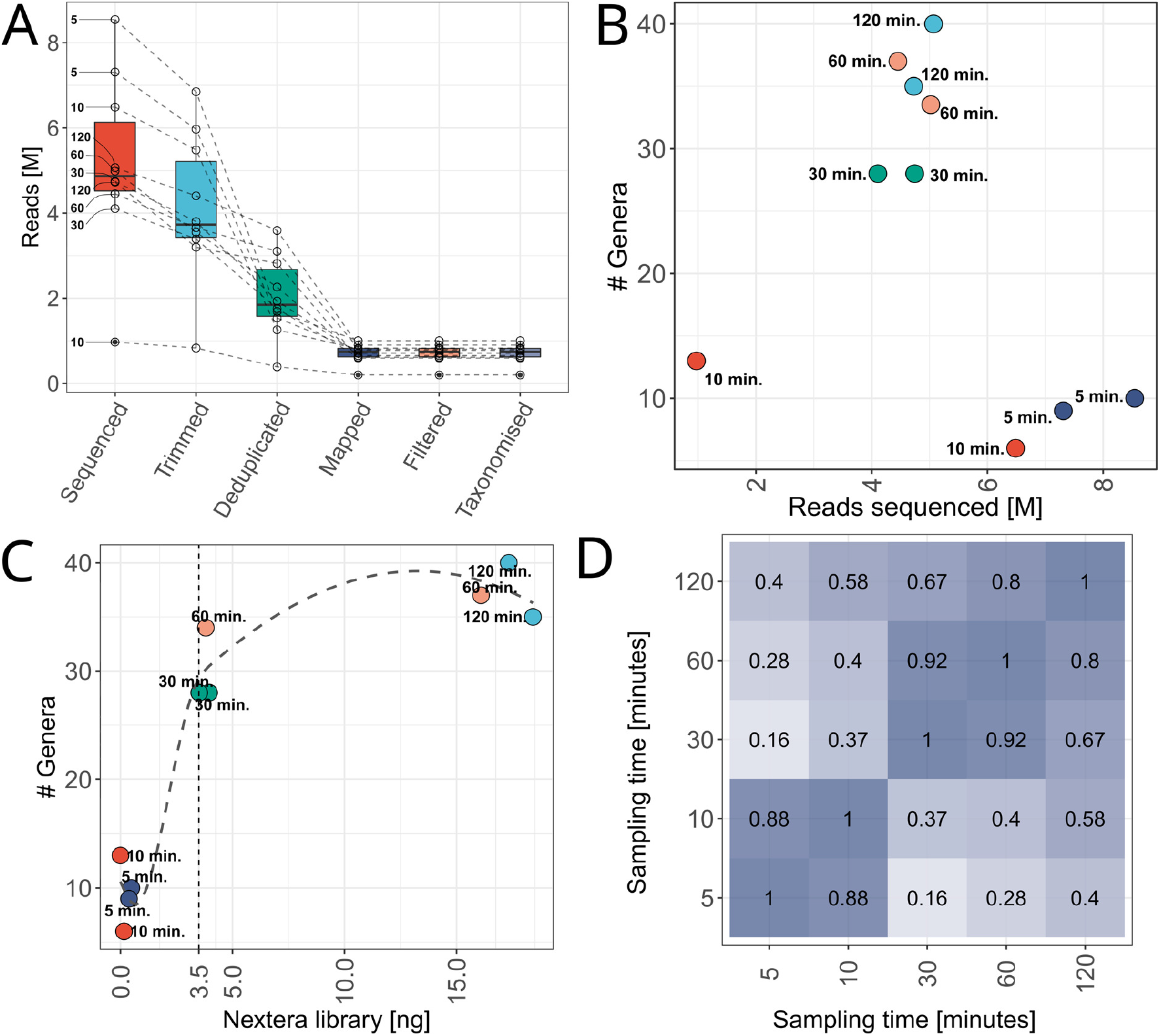
Air-seq performance in a field experiment studying various sampling time. **(A)** Number of reads (million) along the bioinformatic analysis workflow from sequenced to classified reads. The black, bold number next to the ‘Sequenced’ boxplot indicates the sampling time in minutes of a library. Libraries where air was collected for ≤ 10 minutes and that were sequenced to a high number of reads show higher read duplication rates. **(B)** Number of detected genera per million sequenced reads. We observed that a high number of reads in libraries sampled for ≤ 10 minutes does not increase the number of detected genera, whereas in libraries with air-collection times ≥ 30 minutes we observe most genera. **(C)** Number of detected genera compared to the initial library concentration. The longer the sampling time (for our analysed time-points), the higher the final concentration of an Illumina library. The higher the final Illumina library concentration, the more genera we found. We observed the steepest increase in detected genera for Illumina Nextera libraries between 0 - 3.5 ng total yield (smoothed conditional means fitted in R). **(D)** Correlation analysis comparing the RPM per detected genus of all time-points (Pearson correlation coefficients are shown in the boxes). In particular the 30 and 60 minute libraries compared well. One of the 120 minute samples was collected during heavy afternoon rain (but on the same day as all others) potentially affecting correlation analysis to 30 and 60 minute sampling times.

To determine a suitable time-period for future outdoor collections we counted the number of classified genera for each time-point. We found fewest genera at 5 and 10 minutes collection time (10 and 9, as well as 13 and 6 for each sample and time-point respectively) and observed a steep increase in genus numbers from 10 to 30 minutes sampling, with 28 detected genera in both samples at 30 minutes. This number increased even further to 34 and 37 genera in the 60 minute samples and 35 and 40 genera in the 120 minute samples (**Figure 2B, C**). To determine whether the final Illumina library concentration or the sequencing depth contribute to the library complexity (i.e. a higher number of genera in a library) we analysed the number of detected genera per library along with the number of sequenced reads (**Figure 2B**) and the obtained amount of final Illumina library (**Figure 2C**). We found that the nanograms of the constructed Illumina library correlated with the number of observed genera and that libraries containing ≥ 3.5 ng showed the highest number of genera (to obtain a library yield of ≥ 3.5 ng we had to sample air for ≥ 30 minutes at 200 litres per minute) (**Figure 2C**). Although we sequenced some libraries with less than 3.5 ng total yield to a much higher depth than all other samples (**Figure 2B, C**) these samples still contained fewer genera than higher yielding libraries with less sequencing. This indicates that the more metagenomic DNA is present as starting material for the Nextera tagmentation reaction, the higher the likely complexity of a library will be (**Figure 2C**).

We compared the samples using Pearson correlation analysis using the sum of the normalised reads per million (RPM) per detected genus of both samples for each collection time-point. We observed that the 30 and 60 minute sampling time-points correlated best with a Pearson correlation coefficient of 0.92 (**Figure 2D**). For the 30 and 60 to 120 minute samples we observed a drop in the correlation coefficient to 0.67 and 0.80 respectively. One of the 120 minute samples however was affected by heavy afternoon rain potentially inducing a bias in sample composition and so reducing the correlation coefficient.

For future sampling we decided to collect all air samples for 60 minutes: considering that for the 60 minute time-window we found almost as many genera as for 120 minutes sampling, the Coriolis air-sampler can be powered for 60 minutes with one battery load and the observation that the short time (5 and 10 minute) air collection samples suffered from low library concentrations (**Figure 2C**) and higher read duplication rates (**Figure 2A**).

### Analysis of an agricultural field metagenome dynamics over 1.5 months

To assess air metagenome dynamics over multiple weeks in an in-field scenario we collected air for 60 minutes three times a week, in duplicate, between the 12^th^ June and 29^th^ July at the same coordinates (52°37’21.1”N, 1°13’02.6”E) as used for the collection time-series experiment. Samples were collected on Monday, Wednesday and Friday with few exceptions described in **Additional file 2.csv** at the same coordinates as described above. All samples were prepared as Illumina Nextera libraries and sequenced to 5.76 ± 1.88 million 150 base-pair single end reads (**Figure 3A**).

**Figure 3.**
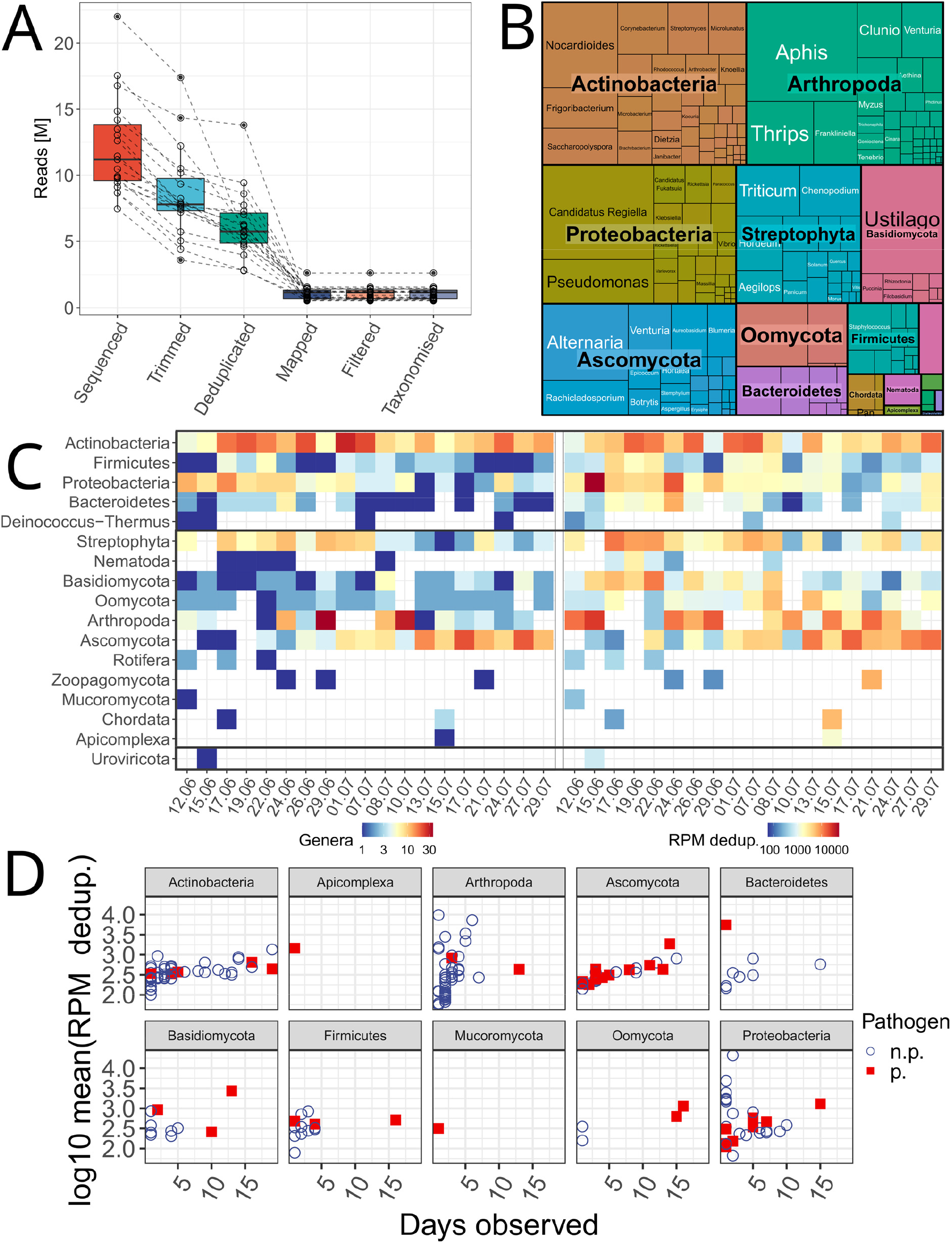
Air-seq experiment at a field site growing wheat, barley and pea crops with air metagenome testing over approximately 1.5 months. **(A)** Number of reads (million) of each sequencing library along the bioinformatic analysis steps undertaken from the number of sequenced reads, to the number of classified reads. **(B)** Abundance (RPM) of all detected phyla over the studied time-period; the larger the box, the higher the abundance of a phylum among all classified reads. The boxes within the coloured phyla boxes show the abundance of the detected genera within a phylum. **(C)** Number of observed genera (left) and RPM counts (right) for the observed phyla. From top to bottom the phyla are grouped in bacteria, eukaryotes and viruses. We observed highest RPM peaks for *Actinobacteria, Proteobacteria, Streptophyta, Arthropoda* and *Ascomycota*. The abundance plots reveal a highly dynamic air metagenome, e.g. with *Streptophyta* displaying higher abundance at the beginning of our sampling series and lesser abundance at later sampling times, an opposite pattern to Ascomycota. **(D)** Phyla with described pathogenic genera filtered using the PHI-base pathogen and pest database [31]. Genera are shown as log10 of the RPM normalised counts over how many days they have been observed by Air-seq. Red squares indicate a genus containing at least one pathogen listed in PHI-base (p.), blue circles a genus which does not have any pathogenic species listed in PHI-base (n.p.). As a particular pathogen rich phylum we identified the *Ascomycota*.

The Coriolis air sampling device was placed in a field immediately surrounded by wheat and barley, as well as peas at a wider distance (**Additional file 1.docx**). We therefore searched the dataset for the presence of the genera *Triticum, Hordeum* and *Pisum*. While we could not detect any *Pisum*, we found *Triticum* as the highest abundant plant genus with 2.23 % of all normalised counts and *Hordeum* as third most abundant plant genus with 1.87 % of all normalised reads immediately after *Chenopodium* (2.04 % of normalised reads) (**Figure 3B and Additional file 3.csv**).

Analysis of read and genus counts over time showed a highly dynamic air metagenome. We detected reads from the eukaryote, bacterial and viral domains. In particular, *Actinobacteria* were present with a high number of genera and with a high number of normalised counts in comparison to other phyla over the entire sampling time-period (**Additional file 3.csv and Additional file 4.csv**). Some phyla showed time specific occurrence patterns: *Arthropoda* were observed in high numbers on specific days. For *Streptophyta* we detected a higher number of genera at the beginning of our sampling series in comparison to later time-points. In contrast, the fungal phylum *Ascomycota* displayed a strong increase of RPM and genera over time (**Figure 3C**).

As many of the detected phyla comprise animal (i.e. and so also human) as well as plant pathogens [31], we analysed our data for the presence of pathogenic genera using the information of the 279 pathogen species listed in the Pathogen Host Interaction database (PHI-base) at phi-base.org [31] (**Additional file 5.csv**). In the phylum *Ascomycota* most genera contain at least one pathogen species known to PHI-base. Considering the noticeable increase in Ascomycota counts in the second half of our sampling time series this potentially indicates the detection of a starting, fungal pathogenic episode by Air-seq (**Figure 3C,D and Additional file 5.csv**).

### Identification and analysis of pathogen species

To identify pathogen species we constructed a smaller database containing a single NCBI RefSeq or GenBank genome per pathogen species of the PHI-base database. We searched this database using BWA-MEM [32], only considering hits mapping with ≥ 95 %-identity and ≥ 95 %-matched sequence exclusively to a single species. We defined an abundance threshold of 0.05 % removing species not supported by 0.05 % of the classified reads within a sample [33].

We found 131 of 271 PHI-base pathogen species in our data targeting a total of 133 host species described in PHI-base. We found that most detected pathogens (96 of 131) colonise plant hosts (**Additional file 6.csv**). We also found 28 pathogens colonising the class mammals, of which 13 also had *Homo sapiens* as a specified host (**Additional file 6.csv**). To test if the detected pathogens colonise the crops grown near our sampling site, we counted the observed pathogen species by plant host genus. Overall we found that the plant genera for which we observed most pathogens comprise major agricultural crops that are actively farmed in Norfolk (**Figure 4A**) [34, 35]. While most pathogens that we detected colonise the genus *Solanaceae* (including e.g. potatoes and tomatoes), *Poaceae* (including wheat and barley) and *Fabaceae* (including peas), which both were growing at our sampling site, were the 2nd and 4th ranked genera with most detected pathogen species along with *Brassicaceae* (e.g. containing cabbages and kale plants) (**Figure 4A**). Specifically, we found 13 wheat, 7 barley and 2 pea pathogen species (**Figure 4B**).

**Figure 4.**
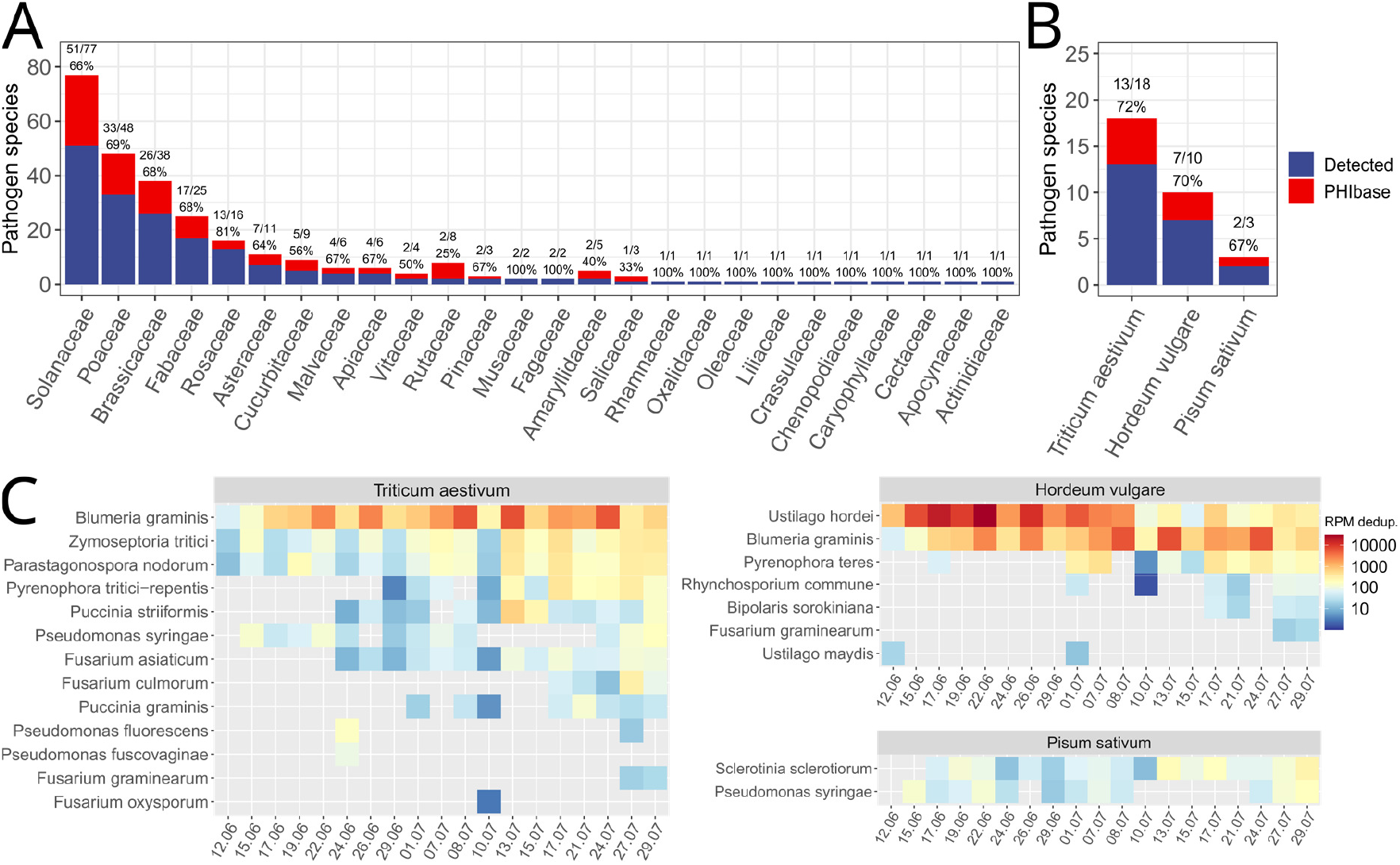
Pests and pathogens detected for various plant genera, wheat, barley and pea. **(A)** The number of detected pathogens and pests for the various plant host families of the PHI-base database. The red bar shows the number of pathogens described in the PHI-base database per host family, the blue bar shows the number of observed pathogens in our Air-seq data. **(B)** The number of pathogens and pests described in the PHI-base database (red) for wheat (*Triticum aestivum*), barley (*Hordeum vulgare*) and pea (*Pisum sativum*) and the number of observed species in our Air-seq data (blue). **(C)** The RPM normalised counts over time of all observed pests and pathogens described to colonise wheat, barley and pea. The species are sorted by the sum of the RPM counts over time from top to bottom. We found *U. hordei* as the most abundant pathogen among the three selected plant species, followed by *B. graminis* as second most abundant wheat and barley pathogen. We detected *U. hordei* with highest abundance at the beginning of our sampling series and fewer counts at the end. Conversely, we detected *B. graminis* with interspersed high RPM peaks over time. In accordance with the above described increase of ascomycete counts at later time-points of our sampling series, we found the ascomycete fungi *B. graminis, Z. tritici, P. nodorum, P. tritici-repentis* and *P. teres* following this pattern with their RPM counts increasing towards the end of our sampling series.

In agreement with our previous finding indicating elevated ascomycete presence at the end of our sampling period, we found especially the ascomycetes *Blumeria graminis, Zymoseptoria tritici, Pyrenophora teres, Parastagonospora nodorum and Pyrenophora tritici-repentis* elevated in our time series, with highest values between 13.07 and 29.07 (**Figure 4B**). At highest counts we found the barley pathogenic basidiomycete *Ustilago hordei* (**Figure 4B**).

As weather influences disease levels [36], we searched for pathogen profiles over time which associate with the weather data. For this we measured the temperature, humidity, rainfall and wind-speed between the 17^th^ of June to the 29^th^ of July at our sampling site. During this time-period we observed three periods of rain and, in particular, elevated humidity in mid-June, mid-July and at the end of July (**Figure 5A**). To match the weather data with pathogen occurrence profiles over time we correlated wheat, barley and pea pathogen RPM values with the weather variables (**Figure 5A**). The occurrence profiles of some pathogens correlated well with the average measured humidity and the average 24 hour rainfall (both important for disease progression [37]) but less with the measured average daily temperature (**Figure 5B**).

**Figure 5.**
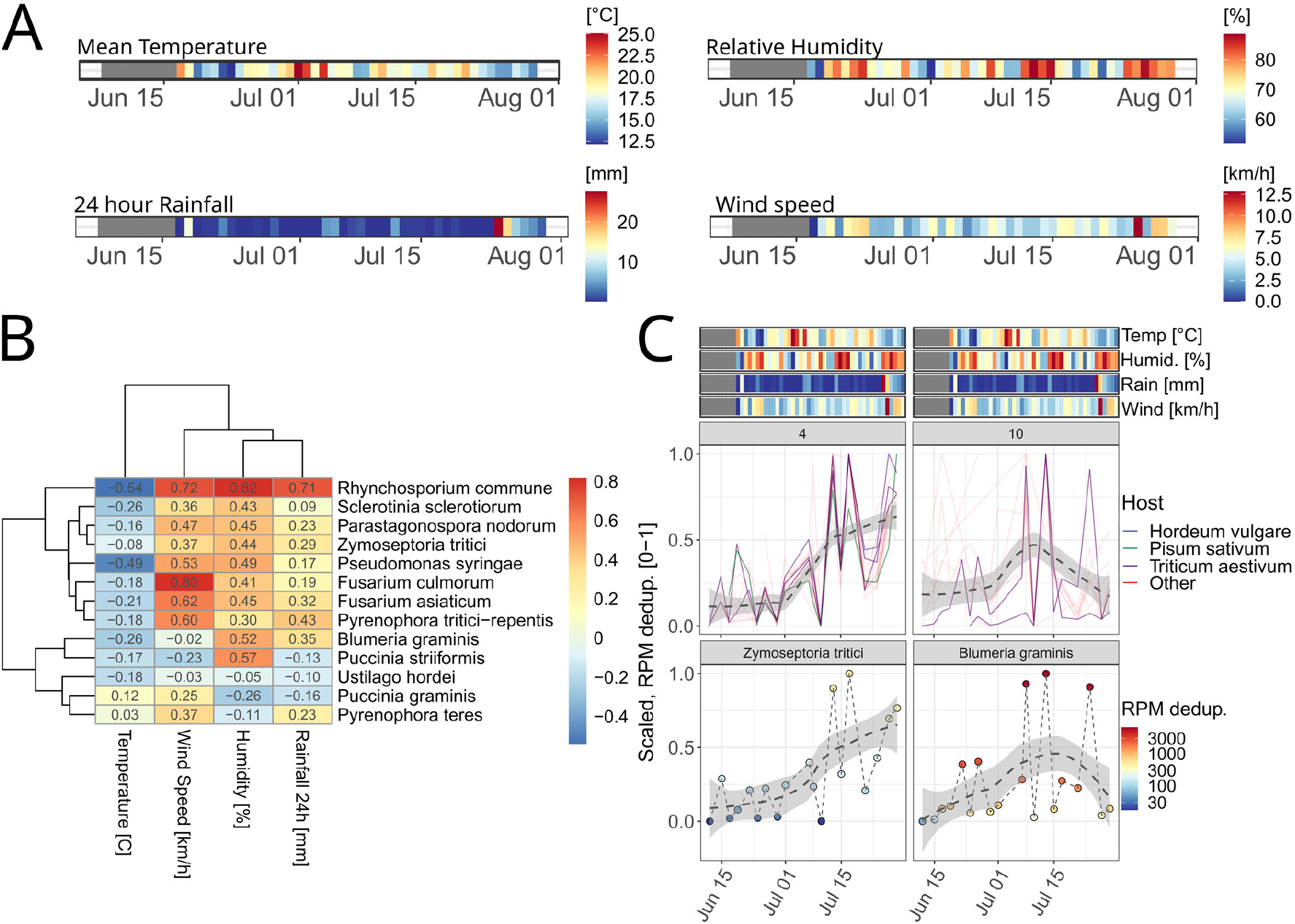
Air-seq pathogen profiles following weather variables. **(A)** Measured weather variables at our sampling site for the air-sampling time-series (grey values indicate non-available weather variables, this data is not available due to a technical fault of the weather station). **(B)** Heatmap showing Pearson correlation coefficients of wheat, barley and pea pathogen RPM values correlated with weather variables. For humidity we observed higher positive correlation values for some pathogens than for other measured weather variables. **(C)** Manually selected clusters that follow measured weather variables well. Top row: For cluster 10 we observed increased RPM counts right after a period of elevated humidity in mid June, mid July and the end of July, for cluster 4 we observed an increase after a period of higher temperatures and elevated humidity in mid July and end of July. Each non-dashed line indicates a species (if coloured, for one of the plant species growing on our sampling site), the grey dashed line shows the smoothed mean over all species in a cluster. The bottom row shows one selected representative species for cluster 4 (*Z. tritici*) and 10 (*B. graminis*).

To visualise and group pathogen levels over time, we clustered the wheat, barley and pea pathogen profiles to similar patterns using the affinity propagation clustering algorithm [38] and so identified 12 clusters (**Additional file 7.csv and Additional file 8.pdf**) for which pathogens showed distinct occurrence profiles For two of the 12 clusters we could detect an increase in pathogen RPM levels at and after the described three periods with rain and elevated humidity levels (**Figure 5C**).

### Assessment of air metagenome pathogen strain information

As whole-genome sequencing in the past has been successfully applied to distinguish between races and strains of plant pathogens [39], we hypothesised that identification of single nucleotide polymorphisms (SNPs) for selected species in our Air-seq data and comparing the variants to a reference database of described, strain specific nucleotide variants will enable us to analyse pathogen dynamics at highest resolution. In 2013 Hubbard et al. established such a reference database by RNA-sequencing and characterising 39 field strains of the obligate fungus and yellow (stripe) rust pathogen *Puccinia striiformis f. sp. tritici* (PST) in and close to East Anglia, a geographical area that includes our sampling site [40]. We compared our Air-seq data for overlaps with SNPs of the 39 field strains. For this we mapped the Air-seq sequencing reads to the *PST-130* EnsemblFungi genome version PST-130_1.0 [41], filtered reads for a %-identity and %-matched-sequence score ? 95 and called SNPs. We excluded all variants with a quality score less than 20 and a read depth less than 20. We also mapped the data of the 39 field strains to the PST-130_1.0 genome using BWA-MEM [32] with default parameters (as described in Hubbard et al.) and called SNPs with the same filtering thresholds as for the Air-seq data. We analysed the transition to transversion ratios of the Air-seq dates with *Puccinia striiformis* calls and the 39 field isolates and found that the values of both datasets agreed well and were slightly larger than 2 (2.65 ± 0.37 for Air-seq and 2.16 ± 0.05 for the PST isolates). Although we did not observe high PST levels in our data (**Figure 4C**), we found a total of 478 SNPs in comparison to the PST-130_1.0 genome after filtering (**Additional file 9.csv**). To find which PST strains were captured by Air-seq we searched for sequence changes that were uniquely assigned to single PST strains among the 478 variants from Air-seq. Of the 478 SNPs we found 20 unique variants of which most were shared with the 13/123 strain that we observed as most abundant on the dates at which we found PST. Other PST strains shared 1 to 4 SNPs with the Air-seq data (**Additional file 10.csv**), potentially indicating that the 13/123 strain or an isolate related to the PST 13/123 strain was captured by Air-seq. The 13/123 strain was collected by Hubbard et al. in Lincolnshire, a county close to our sampling area.

## Discussion

Here we show that airborne shotgun metagenomes of agroecosystems are a rich source of eukaryotic, prokaryotic and viral data and can be utilised for efficient disease monitoring of pests and pathogens at a species and strain level resolution. We do so by collecting air using a battery powered, mobile air sampling device, eDNA extraction and low eDNA input, whole metagenome Illumina sequencing library preparation and analysis. The ability to detect a diverse metagenome in airborne eDNA has been previously reported by Qin et al. [26] and Leung et al. [23] in urban areas. Here we apply shotgun metagenomic profiling of air in a field growing wheat, barley and pea in the UK.

We present a custom workflow enabling researchers to reproduce all steps from collecting air samples to preparing sequencing ready, shotgun metagenome Illumina libraries. We benchmarked our Air-seq workflow assessing its quantitative ability and accuracy. When releasing *B. thuringiensis* spores in a wind tunnel (i.e. a confined environment) we found Air-seq to quantitatively increase *Bacillus* counts with increasing spore release while not detecting any *Bacillus* in the no-spore controls. This agrees well with results obtained by Lynggard et al. [29] and Clare et al. [30] showing the high accuracy of air eDNA profiling methods in zoos - areas with a well described environment.

We found air sampling time to be an important variable determining the genus richness of a sample. In an in-field experiment, we collected air for increasing amounts of time from 5 minutes to 120 minutes consecutively on the same day, observing that longer collections increase the number of observed genera in samples. We successfully prepared and sequenced libraries from 5 and 10 minutes collection time detecting at least 9 and 6 genera in the samples, but found that sampling times ≥ 30 minutes are important to increase observed genus numbers to 28 and more. This agreed with the final library concentrations which also increased with the collection time. Tagmentase enzyme library preparation methods are described to produce higher final library yields with increasing DNA starting amounts [42]. The results therefore indicate that with longer air sampling, more eDNA for library preparation is collected. For our experiments we decided to collect for a consecutive time of 60 minutes and so harvesting 12,000 litres of air at a flow rate of 200 litres per minute using the Coriolis Micro air sampler. We found that this is a convenient time-period using one battery charge of our device at which we were able to detect as many genera as for the 120 minute collection window. The collection time however can vary depending on the environment, the desired application (e.g. rapid air testing or continuous sampling) and on the air-sampling device used. Collection systems from other manufacturers filter air with higher rates than 200 litres per minute and so could reduce collection times (e.g. SASS 4100 Dry Air Sampler with over 4,000 litres per minute; Research International, Monroe, WA, USA).

To study air metagenome dynamics over time we collected air three times a week (Monday, Wednesday and Friday) between the 12^th^ of June and 29^th^ of July in an agricultural environment growing wheat, barley and pea. For this time-period we observed one viral (a double stranded DNA *Lederbergvirus* bacteriophage [43]), 5 bacterial and 11 eukaryotic phyla. As the phyla with highest abundances we found the prokaryotes *Actino*- and *Proteobacteria*, as well as the eukaryotic *Arthropoda, Ascomycota, Basidiomycota* and *Streptophyta*. Especially for eukaryotic phyla we observed temporal dynamics such as a higher number of genera during specific time-periods (e.g. *Streptophyta* at the beginning, *Ascomycota* at the end of our time-series) and varying counts over time (e.g. Arthropoda counts peaking on specific days, *Ascomycota* counts increasing towards the end of our sampling series). Among the characterised genera, we also found the genera of crops growing at our study site at high abundance. We observed 23 plant genera in total with the genus *Hordeum* (includes barley) and *Triticum* (includes wheat) as first and third-most abundant next to the genus *Chenopodium*. The high *Chenopodium* frequency can be explained with *Chenopodium album* being a major weed in wheat and barley crops in temperate and European regions [44, 45] and so contributing to *Chenopodium* presence in the data.

We searched our Air-seq data for pathogen species using the host-pathogen information contained in the PHI-base [31] database. We observed 131 of 271 PHI-base [31] pathogens. Representative for the studied environment we counted 96 of 131 plant pathogens, but we also observed 13 human pathogens. Most pathogens that we detected were described to colonise main crops (e.g. potatoes, tomatoes, wheat, barley, pea) and relatives that are reported to be present in the agricultural area around Norwich [34, 35]. In detail we found 13 wheat, 7 barley and 2 pea pathogen species.

Especially for the grass crops we observed important and devastating species such as the wheat and barley powdery mildew causing *B. graminis* [46] and the wheat Septoria tritici blotch disease causing *Z. tritici* [47]. Powdery mildew (*B. graminis*) and septoria tritici blotch (*Z. tritici*) are among the most common diseases of wheat in the UK. We also detected *U. hordei* as the most abundant barley pathogen. *U. hordei* is the cause of covered smut of barley and common in the UK. Similar to *B. graminis* and *Z. tritici, its* finding at high levels in Air-seq samples therefore can be anticipated and also demonstrates how crops are threatened for a prolonged period for the duration of flowering when they are most susceptible to infection. Among the most abundant wheat pathogens we further found *P. tritici-repentis* and *Parastagonospora nodorum. P. tritici-repentis* and *P. nodorum* respectively, cause tan spot and Septoria nodorum blotch of wheat [48]. The former disease is becoming more common, particularly in warmer, southern parts of the country. The latter disease was once common in the UK but has been very largely replaced by Septoria tritici blotch caused by *Z. tritici*. A PCR-based analysis of leaves of wheat in the UK found up to 100 % of samples to contain DNA of *Z. tritici* with 20 % and 30 %, respectively containing *P. tritici-repentis* or *P. nodorum* of samples to these pathogens [49]. The detection of *P. tritici-repentis* and *P. nodorum* in the Air-seq samples indicates that these pathogens are present on alternative hosts from which spores are released and that could pose a threat to UK wheat crops if environmental conditions are conducive to infection.

We also observed multiple crop pathogenic *Fusarium* species. We detected *Fusarium asiaticum* at highest levels among the detected *Fusarium* species*, which* in its distribution has not been reported in Europe to date [50]. However, genomes of species of the *Fusarium* complex are closely related [50, 51] and when carefully controlling reads assigned to *Fusarium asiaticum* using online BLAST searches, we found equally good hits to multiple *Fusarium* species. This although the BWA-MEM [32] algorithm assigned the 95 %-identity and 95 %-match length filtered reads uniquely to *F. asiaticum* species. It is therefore possible that some of the signals interpreted as coming from these species actually arise from other *Fusarium* isolates that share similarities with *F. asiaticum* [50].

We could relate pathogen frequencies to weather events. We observed multiple fungal crop pathogen species to coincide with elevated humidity and rainfall. This is interesting as the release of ascospores from perithecia is associated with periods of high humidity [52, 53]. We also found several fungal pathogens to coincide well with the measured wind speed. As fungal spores can spread by wind [3, 4], this could indicate the dispersal of spores either from or to our measurement location. Although humidity and wind speeds showed strong positive correlations with pathogen levels, we did not observe similar trends for the measured average daily temperature.

Among the most devastating wheat diseases threatening food production globally are three rusts: *Puccinia graminis f. sp. tritici* (stem rust), *Puccinia triticina* (leaf rust) and *Puccinia striiformis f. sp. tritici* (stripe rust) [54]. Indicating the efficacy of Air-seq in detecting diseases we observed low abundances of *P. striiformis* and *P. graminis* on selected days of our sampling time-series. The detection of *P. graminis* in the Air-seq samples is of particular significance. Stem rust, caused by this pathogen, has been absent from the UK for almost 60 years, but was again observed on a single plant in 2013. Since this time there have been additional reports of stem rust caused by this pathogen [55]. This indicates that the pathogen may not be uncommon and suggests that the relative rarity of stem rust may have been due to environmental factors not being conducive to infection. We also found that some reads mapped to the *P. triticina* genome with a %-identity and %-match score ≥ 95 %, however, at an abundance that was too low to pass the threshold of 0.05 % for classification.

Strain level epidemiology is important to monitor and measure disease threats [40, 56]. While *de novo* identification of strains using metagenomics is challenging, requiring good coverage rates [56], we hypothesised that comparison of nucleotide variants to an established database will allow detailed insights into the presence of pathogen strains at a shallow sequencing depth. Strain profiling using Air-seq methods would therefore enable a scenario where pathogens newly observed in the air can be rapidly evaluated for pathogenicity. A suitable reference dataset for *P. striiformis f. sp. tritici* (PST) strain characterisation has been established in 2013 by Hubbard et al., containing 39 PST UK isolates [40]. The study was performed close to our sampling area. PST is widespread across the western hemisphere and in the past new strains emerged with expanded virulence profiles and higher aggressiveness able to overcome resistance genes in e.g. European germplasm [40, 57, 58]. Hubbard et al. report a shift in UK PST strains with the introduction of a diverse set of new exotic lineages in the last two decades [40].

In a proof-of-concept experiment we mapped and counted variants of our Air-seq reads and the data of Hubbard et al. to the PST-130 reference genome. After quality filtering (%-identity and %-matched-sequence cutoff ≥ 95 for mapped reads, quality threshold > 20 for variant filtering, > 20 depth), we found 20 of 478 variants in our Air-seq data that could be uniquely assigned to a single isolate and so distinguish between PST genotypes. Most of the unique SNPs overlapped with the 13/123 PST strain sampled in Lincolnshire by Hubbard et al. [41], a county close to our sampling site.

Altogether our study adds to the compelling evidence that metagenomic eDNA extracted from air can be used as a source for biomonitoring in agricultural and terrestrial ecosystems [23, 26]. The detection of multiple taxa in air samples tractable to agricultural environments and more importantly the observation of agricultural pathogens colonising the plant species growing at our field site underlines the value of airborne environmental DNA methods. Moreover, the accuracy that we found - with air metagenomics able to detect temporal and weather dependent occurrence patterns as well as to type pathogen species and strains - highlights the value of airborne eDNA in monitoring biodiversity changes and tracking of taxa of human interest or concern.

## Methods

### Air sampling

Air samples were collected using the Coriolis Micro Microbial Air sampler (Bertin Instruments, Montigny le Bretonneux, France) with its tripod fully extended. During sampling we set a flow rate of 200 l/minute for a total of 60 minutes with 10 ml 0.05 % (v/v) Triton X-100 (T8787-50ML, Merck Sigma Aldrich) as collection buffer prepared with UltraPure DNase/RNase free, distilled water (10977023, Invitrogen). After collection we transferred the buffer to a 15 ml conical tube and stored the collected sample at −20 °C until processing. We observed that the collection buffer evaporated during collection to a volume range of 3.5 (on warm days) – 8 ml. To avoid drying of the collection buffer while sampling on especially hot days we topped-up the solution during sampling by briefly stopping air collection, adding 10 ml fresh collection buffer and then continuing with sampling until the 60 minute collection mark was reached (buffer top-up takes approximately one minute). Field samples were collected at a Norwich Research Park field site close to the Earlham Institute (Coordinates: 52°37’21.1”N, 1°13’02.6”E).

### DNA extraction

To standardise the genomic DNA extraction we filtered the collected sample through a 0.22 μm pore size, hydrophilic PVDF, 13 mm diameter filter membrane (GVWP01300, Merck Sigma Aldrich) using a 13 mm diameter stainless steel Swinny filter holder (XX3001200, Merck Sigma Aldrich). We discarded the flowthrough of the PVDF membrane and transferred the membrane immediately after filtration to a 0.7 mm Garnet particle containing tube (13123-50, MoBio – now QIAGEN PowerBead) using sterile tweezers. We immediately added 500 μl UltraPure DNase/RNase free, distilled water to the membrane and ground the reaction for 10 minutes on the TissueLyser II (Qiagen) at 32 Hz. We transferred the supernatant of the grinding step (approximately 300 μl) to a 1.5 ml conical tube and proceeded with DNA purification. DNA purification was performed using the GenFind V2 kit (A41497, Beckman Coulter Life Sciences) according to the standard protocol but with two minor modifications: Our starting volume was 300 μl instead of the 200 μl as in the standard protocol. We therefore adjusted (i.e. increased) the volumes of the buffer solutions as described in the GenFind V2 Blood & Serum protocol (PN B66719AB) for 300 μl starting volume and we reduced the elution buffer volume to 10 μl UltraPure DNase/RNase free, distilled water - in brief: we prepared 450 μl binding buffer (1.5 x sample volume) by combining 440 μl magnetic particle free GenFind V2 Binding Buffer with 10 μl of GenFind V2 Binding Buffer containing magnetic particles. We added the binding buffer to the 300 μl lysate and mixed well. We incubated the reaction for 5 minutes at room temperature, briefly centrifuged the reaction using a table-top centrifuge and pelleted the beads on a magnetic stand. We washed the reaction using 500 μl GenFind V2 Wash Buffer 1, re-pelleted the beads on a magnetic stand and washed the reaction using 500 μl GenFind V2 Wash Buffer 2. We pelleted the beads on a magnetic stand, discarded the supernatant and eluted the extracted DNA in 10 μl water. The extracted DNA was not quantified as DNA amounts were below the detection limit of Qubit 2.0 High Sensitivity reagents (Q32581, Thermo Fisher). All extractions were prepared in a PCR workstation with designated equipment for the workstation. The workstation was UV-light cleaned after each usage.

### Illumina sequencing library preparation

To prepare an Illumina sequencing library from the extracted air DNA we concentrated the 10 μl previously extracted sample to 1.5 μl in an Eppendorf SpeedVac Concentrator 5301. To the 1.5 μl concentrated air DNA extraction we added 2.5 μl Illumina Nextera reaction buffer (FC-121-1030, Illumina), 1 μl 1 pg/μl Lambda DNA (SD0011, ThermoFisher Scientific) as tagmentation reaction control and 0.2 μl Nextera enzyme in a total reaction volume of 5 μl. We incubated the reaction for 5 minutes at 55 °C in a G-Storm GS1 (G-Storm) thermal cycler and after incubation added 5 μl Buffer PB (19066, QIAGEN) to inactivate and strip the transposase from the tagmented DNA. We cleaned the reaction using 10 μl (1 x ratio) AMPure XP beads (Beckman Coulter) and eluted the reaction in 20 μl UltraPure DNase/RNase free, distilled water. We immediately proceeded with library amplification adding 10 μl 5X KAPA 2G Robust Buffer, 1 μl 10 mM dNTPs, 5 μl 2.5 μM P5 and 5 μl P7 oligonucleotide, 8.9 μl water and 0.1 μl KAPA 2G Robust polymerase to the eluted DNA. We incubated the reaction in a G-Storm GS1 thermal cycler with the programme: 3 min at 72 °C, 1 min at 95 °C, [10 s at 95 °C, 30 s at 65 °C, 2 min 30 s at 72 °C] for 18 cycles and a final elongation step for 2 min 30 s at 72 °C. After PCR amplification we cleaned the reaction using 50 μl (1 x ratio) AMPure XP beads eluting in 50 μl 1 x TE buffer. After clean-up we quantified the libraries using Qubit 2.0 High Sensitivity reagents and analysed the size using Agilent Bioanalyzer High Sensitivity DNA Analysis reagents (5067-4626, Agilent) (**Additional file 11.pdf**). As for the DNA extraction, all library preparation reactions were prepared in a designated PCR workstation. The workstation was UV-light cleaned after each usage. All samples were submitted for Illumina HiSeq 2500 150 single-end chemistry sequencing at the Earlham Institute.

### Air metagenome analysis: laboratory background control

Before DNA extraction and library preparation we collected an air sample in our laboratory. Despite working under clean conditions in a PCR working station with assigned pipettes and consumables, we used this sample as background control to consider influences of the lab air environment on our samples. If present, genera found in the lab control sample were routinely removed from the genera detected in all field samples (https://github.com/mgiolai/crop_airseq).

### Air metagenome analysis: NCBI nr database searching

Reads were adaptor and quality trimmed using fastp 0.20.1 [63] and deduplicated using CD-HIT-auxtools 4.6.8 [64]. To analyse the composition of the air samples, we aligned the filtered and deduplicated reads to the entire NCBI nr protein database (downloaded on the 29.10.2021) using DIAMOND 2.0.13 [59] with the settings --evalue 1e-10 --mid-sensitive --max-hsps 1 --top 10 (https://github.com/mgiolai/crop_airseq).

We filtered and classified the diamond output files using custom scripts (https://github.com/mgiolai/crop_airseq) maintaining the DIAMOND e-value threshold ≤ 1e-10 for filtering alignments. Taxonomic binning was performed as described for the MEGAN lowest common ancestor algorithm [33] considering taxonomic assignments of a bitscore within 10 % of the best bitscore. We required taxonomic assignments to be supported by 0.1 % of all classified reads to be valid. Read classification was performed querying the NCBI taxonomy database with the ETE Toolkit 3.1.1 [60] (https://github.com/mgiolai/crop_airseq). The resulting count tables were analysed using custom R-4.1.0 scripts (https://github.com/mgiolai/crop_airseq). Reads were normalised to reads per million by dividing the read number of a taxon with the number of quality filtered and deduplicated reads divided by one million.

### Air metagenome analysis: PHI-base database searching

We downloaded the PHI-base v4.12 pathogen species genomes (one genome per species, 271 of 279 genomes were available, the genomes were queried on 09.12.2022) from NCBI RefSeq and GenBank [61, 62]. We also added the reference genome for *P. triticina* (GCA_019358815.1) to the database. Single genomes were selected as the newest version of the ‘reference genome° or, if a reference genome was not available, as the ‘representative genome° for a species. For multiple available genomes we selected the longest genome by the primary assembly length listed in the NCBI assembly stats file. If neither reference nor representative genomes were available, we selected the longest ‘Complete genome° or if no complete genomes were available the longest genome per pathogen species (https://github.com/mgiolai/crop_airseq). The genomes were combined in a single fasta file. We mapped the fastp 0.20.1 [63] adapter and quality trimmed, as well as CD-HIT-auxtools 4.6.8 [64] deduplicated reads to a database containing a single genome of each pathogen using BWA-MEM 0.7.17 [32] standard settings (https://github.com/mgiolai/crop_airseq).

We filtered the sam files for a %-identity and %-matched sequence score ≥ 95 using filtersam-0.0.11 and analysed the alignments using custom R-4.1.0 scripts (https://github.com/mgiolai/crop_airseq) only considering reads that mapped to a single pathogen species and requiring that a species was supported by 0.05 % of all taxonomically assigned reads. Taxonomic assignment was performed querying the NCBI taxonomy database with the R package taxonomizr 0.8.0 [65]. Reads were normalised to reads per million by dividing the read number of a taxon with the number of quality filtered and deduplicated reads divided by one million.

To cluster the species based on their normalised RPM values (i.e. values scaled from 0 - 1 with 0 as value for no observation and 1 as highest observed RPM value per species) over time, we used the affinity propagation clustering library apcluster [38] in R-4.1.0 (https://github.com/mgiolai/crop_airseq). Correlations with weather values were calculated using the base R-4.1.0 function cor(), selecting pearson as method (https://github.com/mgiolai/crop_airseq).

### Air metagenome analysis: *Puccinia striiformis f. sp. tritici PST-130*

We mapped the fastp 0.20.1 [63] adapter and quality trimmed, as well as CD-HIT-auxtools 4.6.8 [64] deduplicated reads to a database containing a single genome of each pathogen using BWA-MEM 0.7.17 [32] standard settings to the ensembl PST-130_1.0 genome assembly (https://ftp.ensemblgenomes.org/pub/fungi/release-55/fasta/puccinia_striiformis). We filtered the reads for a %-identity and %-matched sequence score ≥ 95 using filtersam-0.0.11 and generated a text pileup from the mapping file using bcftools-1.16 mpileup [66]. We called and filtered variants using bcftools-1.16 [66] with a quality filter setting of -i ‘QUAL > 20 && DP > 20’. Transition to transversion ratios were calculated using the bcftools-1.16 stats command. The variant calling files were analysed using custom R-4.1.0 scripts (https://github.com/mgiolai/crop_airseq).

## Additional Information

**Additional file 1.docx:** Field experiment collection site.

**Additional file 2.csv:** Field experiment collection schedule.

**Additional file 3.csv:** Abundance (% of total) of all detected genera in our field time series.

**Additional file 4.csv:** Observed number of genera per phylum of our field time series.

**Additional file 5.csv:** Genera containing at least one PHI-base pathogen along with number of days observed and mean RPM over time for our field time series.

**Additional file 6.csv:** Detected PHI-base pathogen and corresponding hosts.

**Additional file 7.csv:** All obtained clusters from clustering the normalised pathogen counts over time.

**Additional file 8.pdf:** Images of all obtained clusters from clustering the normalised pathogen counts over time.

**Additional file 9.csv:** Unique SNPs for each of the 39 field isolates sampled by Hubbard et al..

**Additional file 10.csv:** Number of overlapping, unique SNPs for each of the 39 field isolates and the Air-seq data.

**Additional file 11.pdf:** Agilent Bioanalyser High Sensitivity ragent traces of Air-seq Illumina Nextera libraries.

**Additional file 12.txt:** Table with species names, taxonomy ID, genome accession and genome download link of the selected PHI-base v4.12 pathogen genomes and the *P. triticina* genome.

## Data and materials

The datasets generated and analysed during this study are available in the European Nucleotide Archive (ENA) repository under the study accession number PRJEB58191.

## Ethics approval

Not applicable.

## Acknowledgements

The authors acknowledge the Research/Scientific Computing teams at The James Hutton Institute and The National Institute of Agricultural Botany (NIAB) for providing computational resources and technical support for the “UK°s Crop Diversity Bioinformatics HPC” (BBSRC grant BB/S019669/1), use of which has contributed to the results reported within this paper. This research was supported in part by the Norwich Bioscience Institutes (NBI) Computing Infrastructure for Science Group, which provides technical support and maintenance to Earlham Institute’s high-performance computing cluster and storage systems. Next-generation sequencing was delivered at the Earlham Institute, by members of the Platforms and Pipelines team.

## Authors’ contributions

M.G. analysed the data and wrote the manuscript. W.V. established the air-sampling workflow and performed the experiments. R.M.L. and N.P. performed additional data analysis. R.M.L., P.N. and M.D.C consulted on data analysis. W.V., M.D.C. and R.M.L. designed the study. R.M.L. and M.D.C. edited the manuscript.

