## Additional file 1 for "Air-seq: Measuring air metagenomic diversity in an agricultural ecosystem"

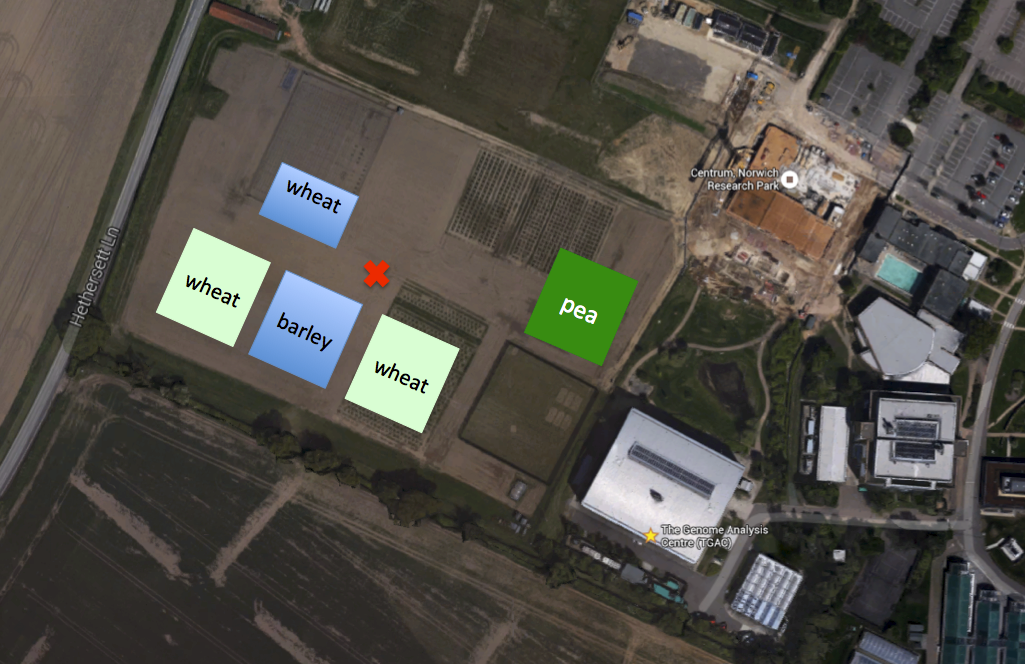

**Figure 1 Sampling location of the Air-seq field experiment:** Air collection was performed three days a week at the highlighted (red cross) spot immediately surrounded by wheat and barley and at further distance pea crops. The Google Earth picture stems from 2013, sampling was performed in 2015.
