## Supplementary figures and images for "Air-seq: Measuring air metagenomic diversity in an agricultural ecosystem"

### Additional file 8

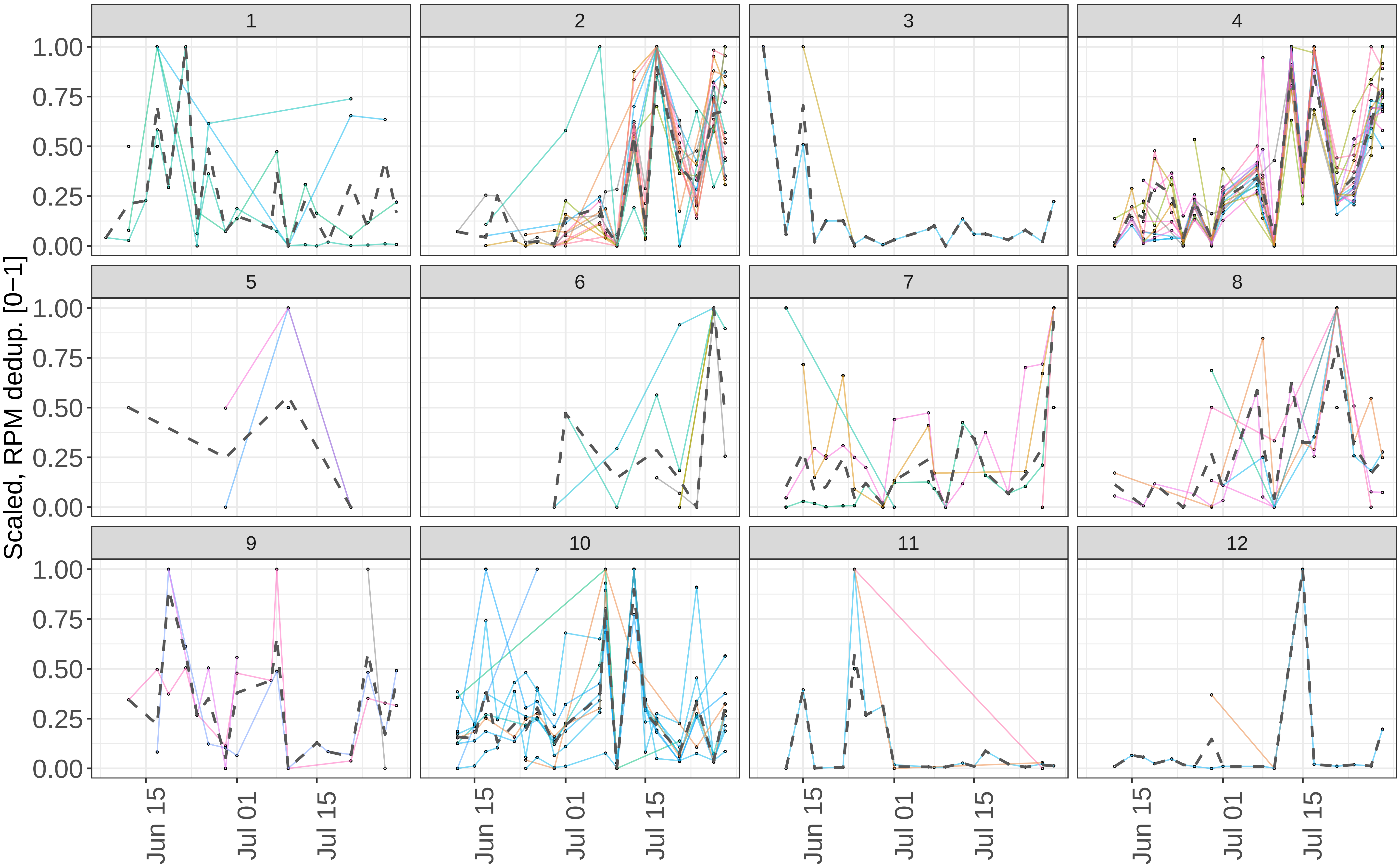
