## Additional file 11 for "Air-seq: Measuring air metagenomic diversity in an agricultural ecosystem"

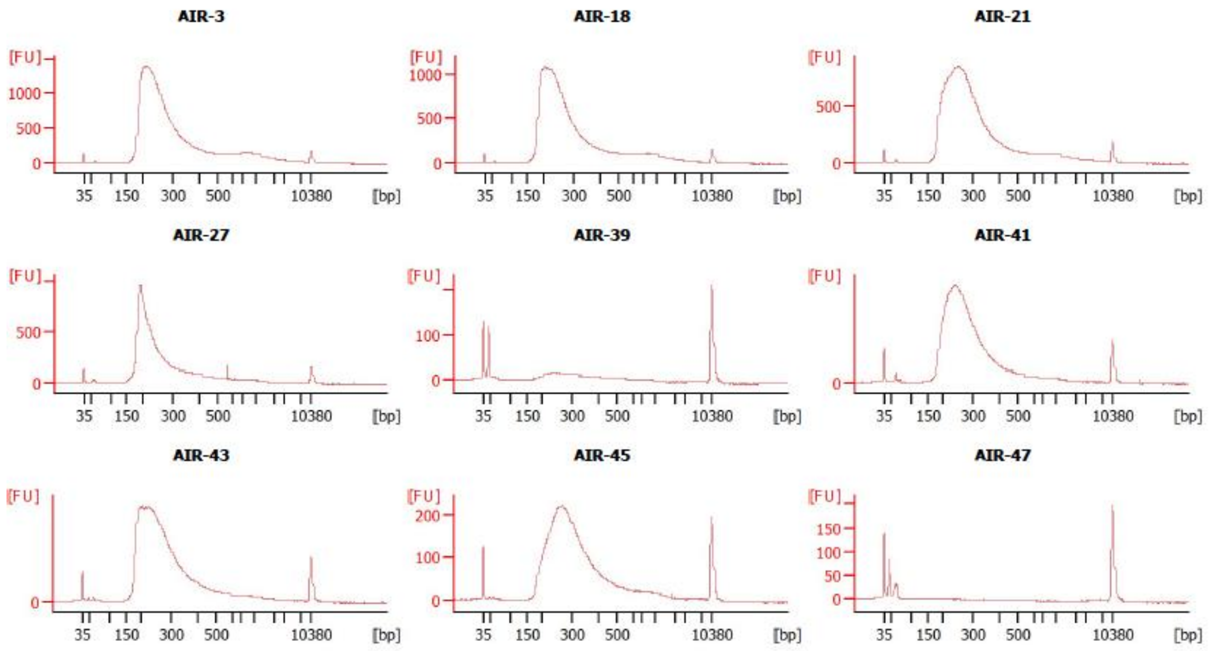


Agilent DNA High Sensitivity reagent traces of final Air-seq Nextera Illumina libraries obtained from samples collected in the field.
